# Histone Deacetylase Inhibitors Prevent Cytokine-Induced β Cell Dysfunction Through Restoration of Stromal Interaction Molecule 1 Expression and Activation of Store-Operated Calcium Entry

**DOI:** 10.1101/2023.12.06.570443

**Authors:** Chih-Chun Lee, Tatsuyoshi Kono, Farooq Syed, Staci A. Weaver, Paul Sohn, Wenting Wu, Garrick Chang, Jing Liu, Marjan Slak Rupnik, Carmella Evans-Molina

**Affiliations:** Center for Diabetes and Metabolic Diseases; Herman B Wells Center for Pediatric Research, Departments of; Pediatrics; Medicine; Anatomy, Cell Biology, and Physiology; Biochemistry and Molecular Biology and; Medical & Molecular Genetics, Indiana University School of Medicine, Indianapolis, IN 46202, USA; Richard L. Roudebush VA Medical Center, Indianapolis, IN 46202, USA; Center for Physiology and Pharmacology, Medical University of Vienna, Austria; Alma Mater Europaea – European Center Maribor, Slovenia; Department of Physics, Indiana University Indianapolis, IN 46202, USA; Department of Physics and Astronomy, Purdue University, West Lafayette, IN 47907, USA

**Author notes:** Correspondence: Carmella Evans-Molina, MD, PhD Indiana University School of Medicine 635 Barnhill Drive MS 2031A Indianapolis, IN. USA.

**Keywords:** Diabetes, pancreatic β cell, store-operated calcium entry (SOCE), stromal interaction molecule 1 (STIM1), calcium, sodium butyrate (NaB)

## Abstract

Histone deacetylase inhibitors (HDIs) modulate β cell function in preclinical models of diabetes; however, the mechanisms underlying these beneficial effects have not been determined. In this study, we investigated the impact of the HDI sodium butyrate (NaB) on β cell function and calcium (Ca^2+^) signaling using ex vivo and in vitro models of diabetes. Our results show that NaB significantly improved glucose-stimulated insulin secretion in islets from human organ donors with type 2 diabetes and in cytokine-treated INS-1 β cells. Consistently, NaB partially rescued glucose-stimulated Ca^2+^ oscillations in mouse islets treated with proinflammatory cytokines. Because the oscillatory phenotype of Ca^2+^ in the β cell is governed by changes in endoplasmic reticulum (ER) Ca^2+^ levels, next we explored the relationship between NaB and store-operated calcium entry (SOCE), a rescue mechanism that acts to refill ER Ca^2+^ levels through STIM1-mediated gating of plasmalemmal Orai channels. We found that NaB treatment preserved basal ER Ca^2+^ levels and restored SOCE in IL-1β-treated INS-1 cells. Furthermore, we linked these changes with the restoration of STIM1 levels in cytokine-treated INS-1 cells and mouse islets, and we found that NaB treatment was sufficient to prevent β cell death in response to IL-1β treatment. Mechanistically, NaB counteracted cytokine-mediated reductions in phosphorylation levels of key signaling molecules, including AKT, ERK1/2, glycogen synthase kinase-3α (GSK-3α), and GSK-3β. Taken together, these data support a model whereby HDI treatment promotes β cell function and Ca^2+^ homeostasis under proinflammatory conditions through STIM1-mediated control of SOCE and AKT-mediated inhibition of GSK-3.

## Introduction

Diabetes, a chronic condition that affects more than 34 million people in the United States, is caused by near complete loss of insulin secretion in the case of type 1 diabetes (T1D) or insulin resistance that is accompanied by inadequate insulin secretion in type 2 diabetes (T2D) (1). Altered β cell endoplasmic reticulum (ER) Ca^2+^ signaling has been linked with the development of both T1D and T2D and leads to reduced insulin secretion, activation of ER stress, and β cell death (2). Under steady state conditions, the sarco-ER Ca^2+^ ATPase (SERCA) pump transports Ca^2+^ into the ER lumen, while inositol triphosphate (IP_3_) and ryanodine receptors release Ca^2+^from the ER (3). When ER Ca^2+^ levels are depleted, a tightly regulated mechanism known as store-operated calcium entry (SOCE) is activated by the ER Ca^2+^ sensor stromal interaction molecule 1 (STIM1) to refill Ca^2+^ stores through gating of plasmalemmal Orai channels (4).

Pharmacological inhibition of SOCE alters glucose-stimulated Ca^2+^ oscillations and reduces insulin secretion in murine pancreatic islets and clonal β cell lines (5-7). Moreover, STIM1 gene deletion in mice leads to the loss of SOCE in islets and decreases glucose-stimulated insulin secretion (GSIS) that is mediated through the GPR40 agonist fasiglifam, a modulator of IP_3_ production and Ca^2+^ release (7). Notably, lower STIM1 expression has been observed in human islets from donors with T2D and in mouse islets and INS-1 832/13 pancreatic β cells (INS-1 cells) treated with proinflammatory cytokines and free fatty acids, showing that reduced insulin secretion, ER Ca^2+^ depletion, and increased β cell ER stress are associated with lower STIM1 levels (6). Conversely, STIM1 overexpression increases insulin secretion in human islets isolated from organ donors with T2D (6). However, at present, the pathways leading to altered SOCE in the diabetic β cell remain largely unexplored.

Histone deacetylases (HDACs) control various cellular processes by removing acetyl and acyl groups from lysine residues of histone and non-histone proteins. HDACs play a central role in the epigenetic regulation of gene transcription: histone lysine deacetylation increases chromatin compaction, reducing the accessibility of the transcriptional machinery to gene targets and, therefore, suppressing gene transcription. More recently, HDACs have been shown to regulate pathways outside of the nucleus through the acetylation of non-histone proteins (8).

Notably, HDAC inhibitors (HDIs) have been linked to a number of beneficial metabolic effects, including protection against diet-induced obesity in mice, potentiation of GSIS through enhanced GLP-1R agonism, and protection of pancreatic β cells from cytokine-induced apoptosis (9, 10). However, the mechanisms by which HDIs have these pleiotropic effects are not fully understood. Herein, we highlight a novel pathway through which HDAC inhibition exerts protective metabolic effects in the β cell through upregulation of STIM1 and restoration of SOCE. Our results reveal a unique ability of NaB to rescue IL-1β-mediated reductions in SOCE, suggesting a previously unrecognized relationship between SOCE and HDI that could impact future therapeutic strategies to reduce the deleterious effects of inflammatory states in diabetes.

## Materials and Methods

### 2.1 Materials

Sodium butyrate (NaB), diazoxide, verapamil, and recombinant human interleukin-1β (IL-β) were obtained from Millipore Sigma (St Louis, MO, USA). Thapsigargin (TG) was purchased from Cayman Chemicals (Ann Arbor, MI, USA). The mouse interferon-γ (IFN-γ) and human tumor necrosis factor-α (TNF-α) used in the incubation of pancreatic β cells and isolated islets were purchased from Invitrogen (Carlsbad, CA, USA). Ca^2+^ imaging of mouse islets was performed with Fura-2 AM (Fura-2-acetoxymethyl ester) acquired from Thermo Fisher Scientific (Waltham, MA, USA).

### 2.2 Animals, islets, and cell culture

C57BL/6J mice (Jackson Laboratories, Bar Harbor, ME, USA) were maintained under a protocol approved by the Indiana University Institutional Animal Care and Use Committee (IACUC). Mice were kept on a standard light-dark cycle with *ad libitum* access to food and water. Mouse pancreatic islets were isolated by collagenase digestion, hand-picked, and allowed to recover overnight as described previously (11). Rat INS-1 832/13 (INS-1) cells (RRID:CVCL_7226) were cultured as previously described (12). STIM1 knockout (STIM1 KO) INS-1 cells were generated by CRISPR/Cas9 genomic editing techniques as described in (6). Cadaveric human islets from donors with T2D were obtained from the Integrated Islet Distribution Program (IIDP). Upon receipt, human islets were hand-picked and allowed to recover overnight in DMEM supplemented with 5.5 mM glucose, 10% fetal bovine serum, and 100 units/mL penicillin-streptomycin. Data presented include two donors with T2D: one male donor, 63 years old, BMI 31.7 kg/m^2^; and one female donor 65 years old, BMI 30.4 kg/m^2^.

### 2.3 Glucose-stimulated insulin secretion

Glucose-stimulated insulin secretion (GSIS) assays were performed as previously described (6). Briefly, cultured INS-1 cells or isolated islets were incubated for 2 hours in 1x secretion assay buffer (SAB) containing 2.5 mM glucose and then in 1x SAB containing 15 mM glucose for an additional 2 hours. Insulin secretion was measured in the harvested liquid using either a human insulin ELISA kit (Cat#10-1132-01, Mercodia, Winston Salem, NC, USA) or a rat insulin ELISA kit (Cat#10-1250-01, Mercodia, Winston Salem, NC, USA) according to the manufacturer’s instructions. The secreted insulin levels were normalized to total protein content.

### 2.4 Ca^2+^ imaging

Intracellular Ca^2+^ was measured using the FLIPR Ca^2+^ 6 Assay Kit and a FlexStation 3 system (Molecular Devices, Sunnyvale, CA, USA). In brief, INS-1 cells were plated in black wall/clear bottom 96-multiwell plates (Costar, Tewksbury, MA, USA), cultured to ∼90% confluency, and treated with drugs and/or stress conditions as indicated. Following the specified treatment, calcium 6 reagent was added to the medium, and cells were incubated for an additional 2 hours at 37 °C and 5% CO_2_. Cells were then incubated for ∼4 minutes in Ca^2+^-free Hanks’ balanced salt solution (HBSS) containing 5.5 mM glucose, 0.5 mM EGTA, 10 µM verapamil, and 200 µM diazoxide. The baseline fluorescence (F0) was measured for a minimum of 10 seconds in a Ca^2+^-free HBSS medium. TG was then added at ∼10 seconds to deplete ER Ca^2+^ stores, followed by supplementation with Ca^2+^ at ∼140 seconds to reach 2 mM Ca^2+^ in the medium. SOCE was determined by the formula ΔF/F0, where ΔF is the increase in calcium 6 fluorescence intensity. Data were acquired at 1.52-second intervals using an excitation wavelength of 485 nm and an emission wavelength of 525 nm.

Imaging of glucose-stimulated Ca^2+^ oscillations in islets was performed using islets from mice expressing GCaMP6s, which is a genetically encoded fluorescent indicator of cytosolic Ca^2+^. The GCaMP6s expressing mice were obtained by crossing *Ins1*^Cre^ mice (The Jackson Laboratory, Strain #26801) with GCaMP6s Cre-inducible mice (The Jackson Laboratory, Strain #028866). Isolated mouse islets were allowed to recover overnight and then treated for 24 hours with NaB and/or a cytokine mixture containing 5 ng/mL IL-1β, 100 ng/mL IFN-γ, and 10 ng/mL TNF-α. Baseline measurements were performed at 5.5 mM glucose and Ca^2+^ transients were measured in response to 11 mM glucose using a Zeiss Z1 microscope as described previously (6). Analysis of Ca^2+^ imaging data and activity was performed according to previously described methods and quantitated as an activity under basal 5.5 mM glucose conditions and during the 1^st^ phase responses to glucose (13, 14). In brief, regions of interest (ROIs) were identified using automatic detection. Significant changes in cytosolic Ca^2+^ were automatically annotated as events (with z score > 3) within the ROIs. Events were characterized by their start time, peak time, and the width at half of their peak amplitude (events halfwidth). Activity at each of the indicated phases was quantitated as the sum of AUC of individual detected events; sums were log_10_ transformed for presentation.

The analysis of the Ca^2+^ oscillatory signal necessitates an understanding of the distinct frequencies present within the time series data. In brief, the oscillatory time series data underwent initial processing using singular spectrum analysis (SSA). This technique decomposes the time series data into a serial of singular functions with multiple eigen-frequencies, each function possessing a corresponding amplitude coefficient that signifies its contribution to the original time series (15). This decomposition method facilitates the segregation of oscillation data into components with varying eigen-frequencies, as well as minor fluctuations and overall trends. Components exhibiting lesser contributions were identified as noise and consequently excluded from subsequent analysis. Subsequently, the Fourier transform was employed to convert the components primarily responsible for the oscillatory signals from the time domain to the frequency domain (16). A peak detection algorithm was applied to the Fourier spectra to identify the significant frequencies within the oscillations. Finally, violin plots were generated to visualize the prevalent significant frequencies. We used Jenks optimization method to determine the different populations for all detected frequencies and the separation breaks between those populations. Two groups of oscillation frequencies were identified: the slow base oscillation and the fast oscillation. The threshold of separation between these two groups can be used to calculate the fast Ca^2+^ oscillation frequency percentage for each islet condition.

### 2.5 Cell viability assay

INS-1 cells were cultured in black wall/clear bottom 96-multiwell plates to reach ∼70% confluence. Following treatment with NaB and/or IL-1β for 24 hours, 100 μL of CellTiter-Glo Luminescent Cell Viability Assay reagent (Promega, Madison, WI, USA) was added to the medium. After a 10-min incubation at room temperature, luminescence was measured with a plate reader (SpectraMax iD5 Multi-Mode microplate reader, Molecular Devices). The luminescent reading of vehicle-treated cells was set as 100%, and the percentage viability relative to vehicle groups was reported.

### 2.6 Immunoblot

Immunoblot experiments were performed as previously described (17, 18) using antibodies against cleaved caspase-3 (Cell Signaling Technology, Cat#9664, RRID: AB_2070042), procaspase-3 (Cell Signaling Technology, Cat#9662, RRID: AB_331439), STIM1 (BD Biosciences, Cat#610954, RRID: AB_398267), and β-actin (Millipore Sigma, Cat#MAB1501, RRID: AB_2223041). All primary antibodies were used at a dilution of 1:1000. Species-matched secondary antibodies (LI-COR Bioscience, Cat# 926-32212; RRID: AB_621847 and LI-COR Bioscience, Cat# 926-68073; RRID: AB_10954442) were used. The images were analyzed using LI-COR Biosciences Image Studio.

### 2.7 Quantitative RT-PCR

Total RNA was extracted from INS-1 cells or isolated islets using an RNeasy Mini Plus kit or RNeasy Micro Plus kit, respectively (Qiagen, Valencia, CA, USA). Isolated RNA was reverse-transcribed into cDNA using M-MLV Reverse Transcriptase (Invitrogen, Carlsbad, CA, USA) as described previously (19). The cDNA was subjected to quantitative PCR analysis using SsoAdvanced Universal SYBR Green Supermix (BioRad, Hercules, CA, USA). Relative gene expression levels were established against the housekeeping gene β-actin, using the comparative delta Ct method, as described previously (20). Primer sequences are provided in Table 1.

**Table 1:**
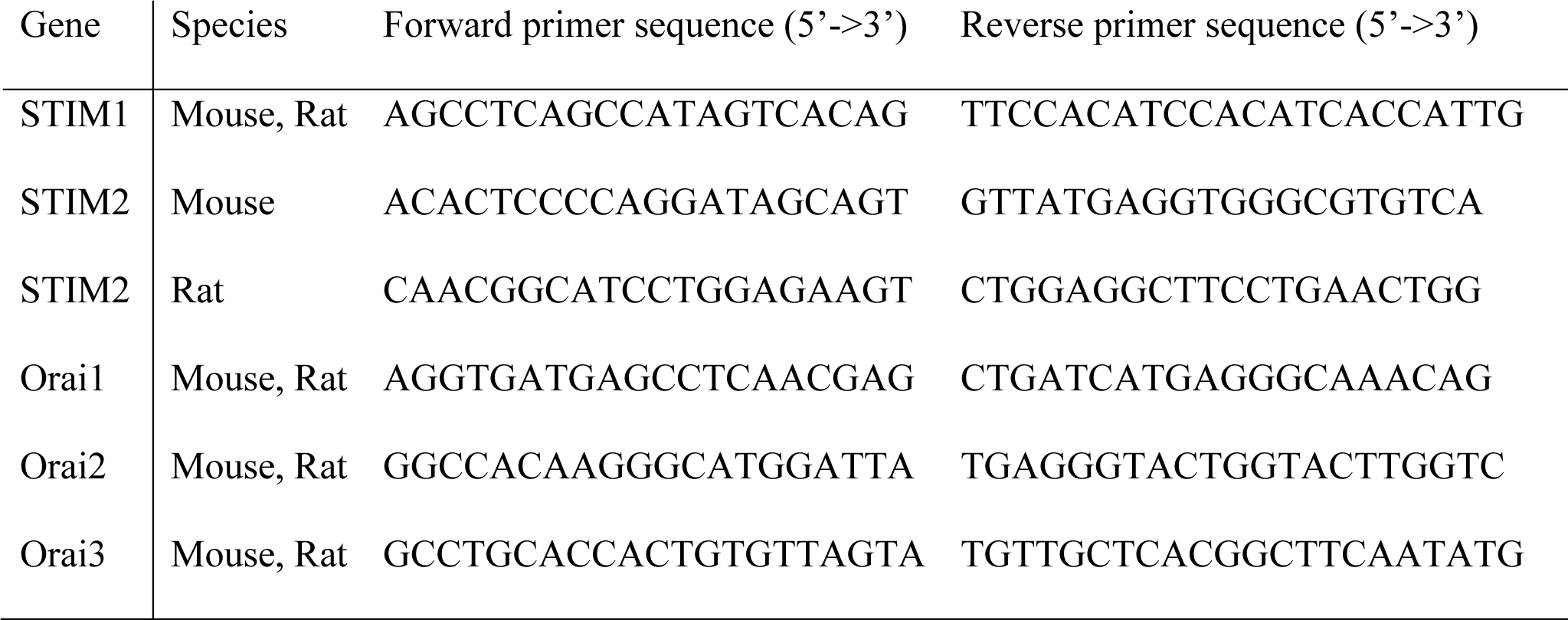
List of primers used in the study.

### 2.8 Phosphorylation array assay

C57BL/6J mouse islets were isolated and allowed to recover overnight. Islets obtained from 14 mice were pooled and ∼700 hand-picked islets were distributed for each treatment group. The islets were treated with NaB and/or cytokines (IL-1β, IFN-γ, and TNF-α) for 16 hours. The treated islets were lysed using the lysis reagent provided in the phosphorylation array kit from RayBiotech Life (Peachtree Corners, GA, USA, Cat#AAH-MAPK-1-8). The lysates were incubated with membranes pre-coated with anti-phosphorylated proteins overnight and incubated with anti-rabbit IgG (detection antibody cocktail) overnight. The membranes were then incubated with HRP-conjugated anti-IgG. The membranes were incubated in a detection buffer and imaged with a chemiluminescent imaging system. Fiji software was used to analyze the resulting data.

### 2.9 Statistical analysis

Unless otherwise indicated, results are presented as the mean ± SEM. The statistical significance of differences between groups was analyzed using GraphPad Prism software (San Diego, CA, USA). Comparisons between the two groups were performed using an unpaired student’s t-test, while comparisons among multiple groups used one-way ANOVA with Tukey-Kramer post-hoc analysis. A *P*-value < 0.05 was considered statistically significant.

## Results

### 3.1 Inhibition of HDAC increases insulin secretion in human islets from donors with T2D and INS-1 cells treated with IL-1β

To test whether NaB has beneficial effects on insulin secretion in humans, islets isolated from human cadaveric donors with T2D were incubated in the absence or presence of NaB for 24 hours prior to a GSIS assay. NaB treatment significantly increased insulin secretion in islets from T2D donors in comparison to vehicle-treated T2D islets (Figure 1A). To explore whether NaB had similar effects under pro-inflammatory conditions that typify T2D (21), we used INS-1 cells treated with IL-1β. IL-1β treatment abolished GSIS in INS-1 cells, yet this secretory deficit was partially prevented when cells were co-incubated with IL-1β and NaB (Figure 1B).

**FIGURE 1.**
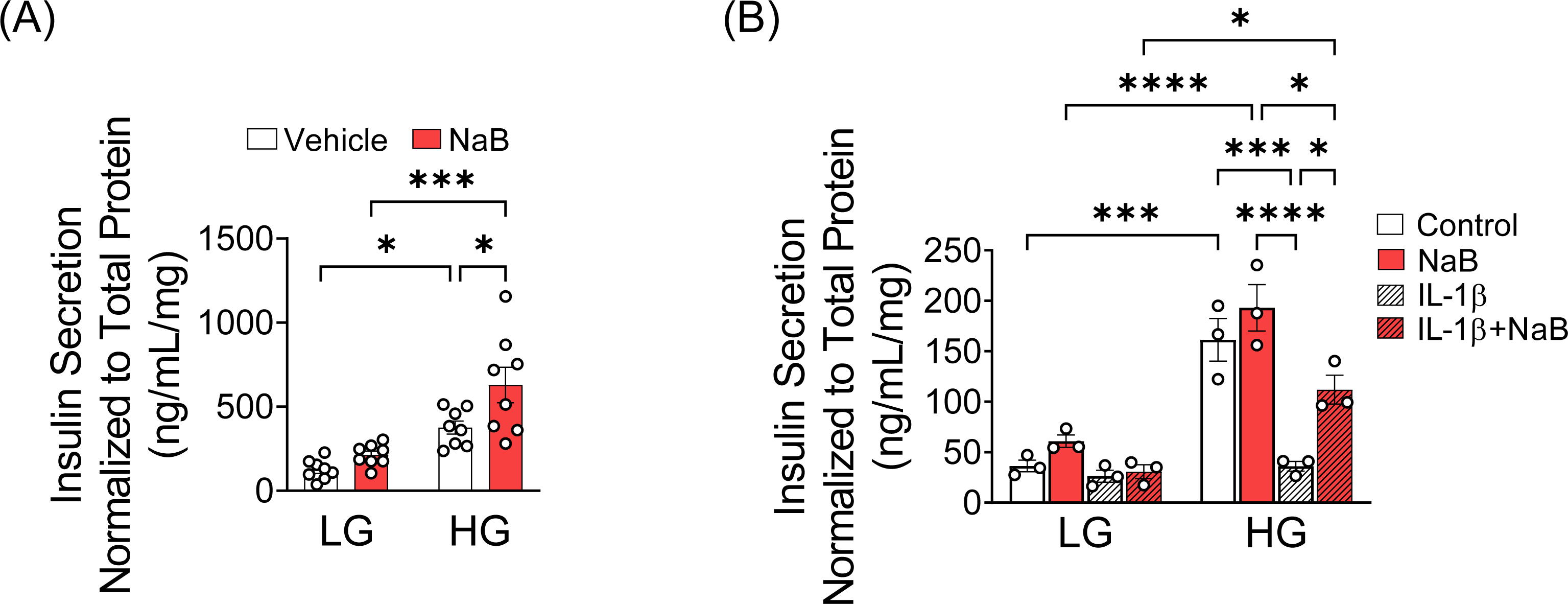
Inhibition of HDAC increases insulin secretion in human islets from donors with T2D and INS-1 cells treated with IL-1β. (**A**) Human islets from two donors with type 2 diabetes (T2D) were incubated with or without 0.5 mM NaB for 24 hours, and GSIS was performed. Insulin levels were determined using a human insulin ELISA kit and normalized to total protein content in the lysates. The open circles indicate ELISA replicates from 2 donors. (**B**) INS-1 cells were treated with IL-1β and GSIS was performed. Insulin levels were measured using a rat insulin ELISA kit and data were normalized to total protein content. The values obtained in 3-4 independent experiments are represented by open circles. Results are shown as the mean ± S.E.M; \**P* < .05, \*\*\**P* < .001, \*\*\*\**P* < .0001.

### 3.2 Inhibition of HDAC prevents cytokine-induced dysfunction in Ca^2+^ oscillations

Impaired insulin secretion is associated with defective Ca^2+^ homeostasis and impaired stimulus-secretion coupling (22), and we have previously linked changes in Ca^2+^ oscillations with alterations in glucose-stimulated Ca^2+^ responses in islets (6). Therefore, to test whether NaB impacts Ca^2+^ homeostasis, islets were isolated from *Ins-1* cre-dependent GCaMP6s mice that express a fluorescent indicator specifically in β cells (23). The isolated islets were treated for 24 hours with a combination of IL-1β, IFN-γ, and TNF-α in the presence or absence of NaB, and glucose-stimulated Ca^2+^ oscillations were measured. In the absence of cytokine treatment, vehicle and NaB-treated islets showed apparent first-phase responses followed by second phase Ca^2+^ oscillations in response to glucose stimulation, as previously reported (6) (Figure 2A-B). However, upon cytokine treatment, the first-phase Ca^2+^ response was significantly reduced while it was restored by NaB treatment (Figure 2C-E). Quantitative analysis of the fraction of active β cells during glucose-dependent activation also showed that cytokine treatment resulted in loss of the first phase glucose response, which was restored by NaB treatment (Figure 2F). To determine the effect of NaB on second phase responses, frequencies from the Ca^2+^ oscillations from each group were extracted and represented in violin plots (Figure 2G). All violin plots exhibited a comparable distribution in the lower frequency range, suggesting a fundamental slow oscillatory signal common to all conditions. However, a significant number of frequencies detected from cytokine-treated islets (32.09%) were in the fast oscillation (high frequency) range, as compared to the low percentage in this range seen in vehicle-treated and NaB-treated islets (5.54% and 1.21%, respectively) (Figure 2G). We also observed that the accelerated frequencies in the second phase Ca^2+^ oscillations due to cytokine stress were drastically diminished by co-treatment with NaB (1.56% of detected frequencies were fast oscillations). These data suggest that NaB treatment suppresses cytokine-dependent effects on both first phase and second phase Ca^2+^ responses in islets.

**FIGURE 2.**
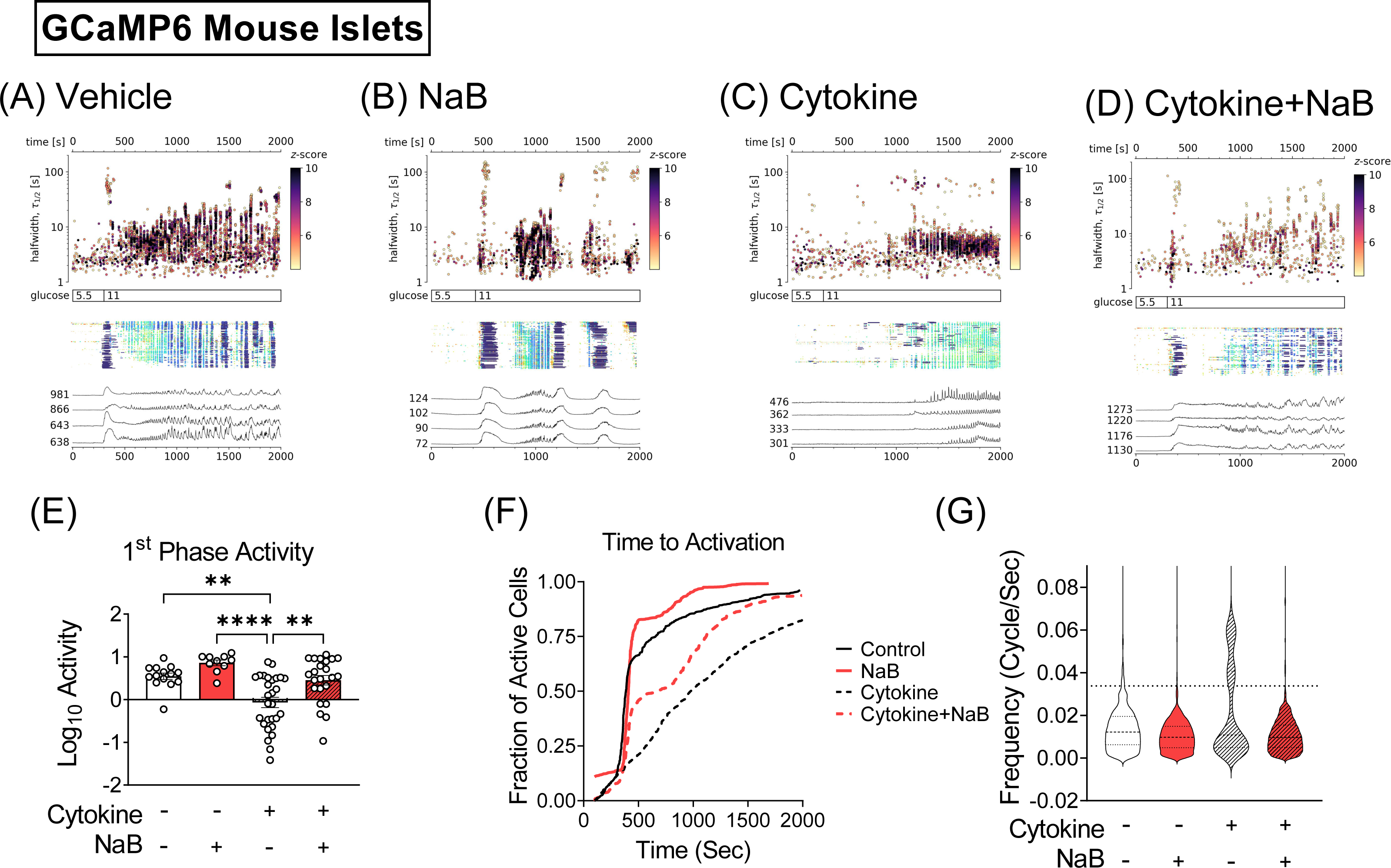
Inhibition of HDAC partially prevents cytokine-induced dysfunction in Ca^2+^ oscillations. Islets obtained from mice carrying the Ins-Cre inducible GCaMP6s Ca^2+^-binding fluorescent indicator were treated with 5 ng/mL IL-1β, 100 ng/mL IFN-γ, and 10 ng/mL TNF-α in the absence or presence of 0.5 mM NaB for 24 hours. The number of islets measured in each group was as follows: vehicle (n=14),), NaB (n=11), cytokine (n=27), and cytokine + NaB (n=14). Individual cells were determined as regions of interest (ROIs) obtained by our automated segmentation algorithm. We discarded ROIs with number of events below 5. (**A-D**) Top panels: [Ca²⁺] events’ halfwidth duration through time. Color indicates the statistical significance in terms of z-score. Middle panels: A raster plot of cytoplasmic Ca²⁺ concentration showing events in individual ROIs. Color indicates the duration of events, with long events indicated in blue and short events in red. Lower panels: Representative cytosolic Ca^2+^ traces after treatment with 11 mM glucose from a number of ROIs that represent best the average activity of the islets. (**E-F**) Activity for the baseline and phase 1 and phase 2 responses is displayed as the log10 value. Results are displayed as means ± SEM; *p<0.05, **p<0.01, ****p<0.0001. (**G**) Pooled data showing the fraction of active cells during glucose-dependent activation. (**H**) Violin plots of the significant frequencies detected from the Fourier transform of singular spectrum analysis (SSA) components. Dashed line indicates the cluster separation threshold determined by Jenks optimization method.

### 3.3 Inhibition of HDAC improves SOCE in IL-β-treated INS-1 cells

Our previous studies showed that changes in Ca^2+^ oscillatory activity were associated with impaired SOCE (6). Therefore, to determine if NaB impacts cellular Ca^2+^ homeostasis and SOCE under cytokine stress, INS-1 cells were treated for 24 hours with IL-1β in the absence or presence of NaB, and the depletion of ER Ca^2+^ and subsequent SOCE-mediated replenishment of Ca^2+^ stores was characterized (Figure 3A). First, the uptake of Ca^2+^ from the extracellular milieu was eliminated by blockade of plasma membrane Ca^2+^ channels with diazoxide and verapamil and incubation of the cells in Ca^2+^ free buffer with EGTA chelation. The first measured peak of cytosolic Ca^2+^ reflects the release of stored Ca^2+^ from the ER via dosing with TG, which inhibits SERCA-mediated Ca^2+^ uptake to the ER. Next, extracellular Ca^2+^ was supplemented, allowing Orai- and STIM1-mediated SOCE, which was measured in the second peak of cytosolic Ca^2+^. As shown previously (6), treatment with IL-1β significantly reduced ER Ca^2+^ stores (a negligible first peak in response to TG) and SOCE (reduced second peak following Ca^2+^ supplementation) (Figure 3B-E). In contrast, simultaneous incubation of NaB and IL-1β partially maintained ER Ca^2+^ stores (Figure 3B-D) and SOCE responses (Figure 3B-C and E) in INS-1 cells.

**FIGURE 3.**
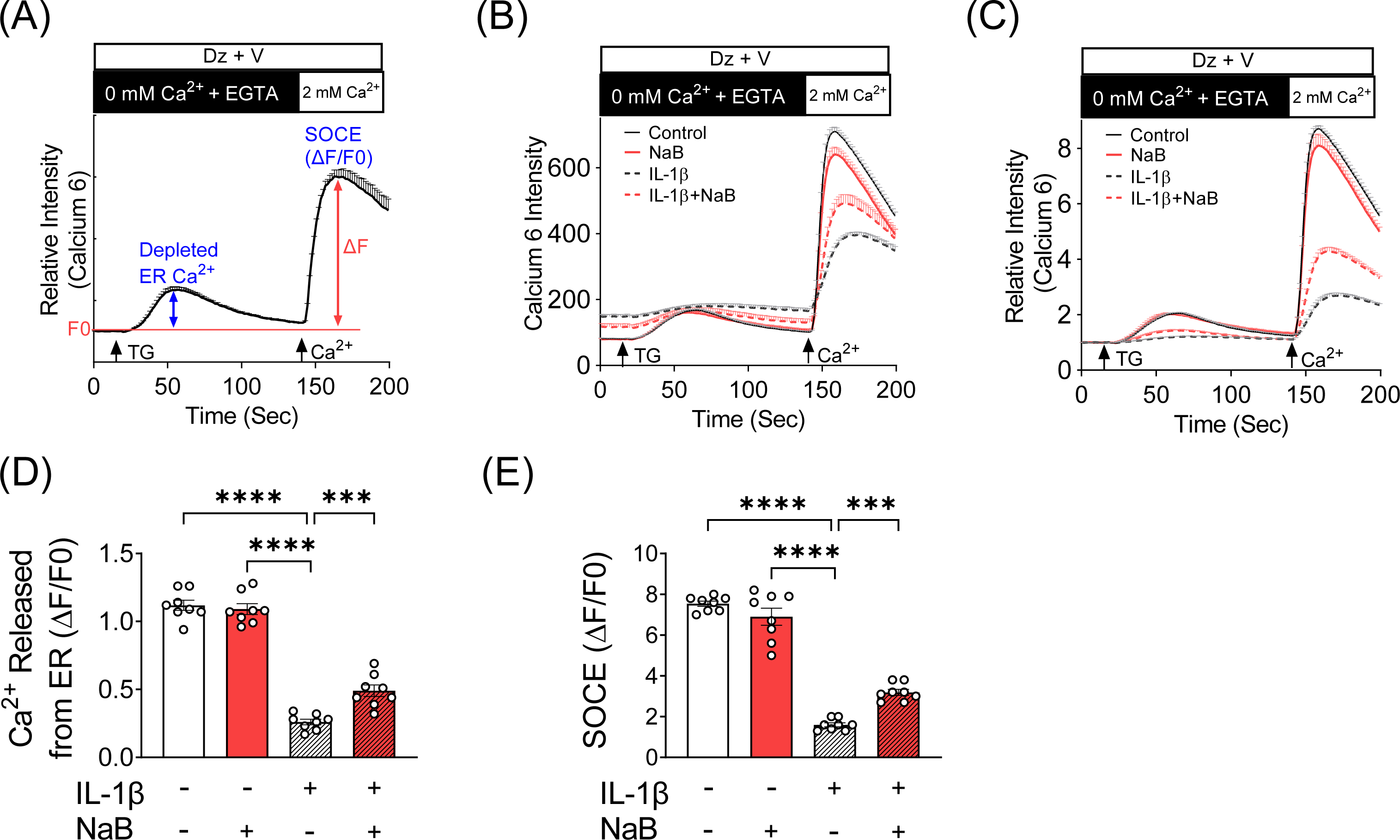
Inhibition of HDAC improves SOCE in IL-β-treated INS-1 cells. (**A-E**) Levels of stored ER Ca^2+^ and store-operated Ca^2+^ entry (SOCE) were measured with Calcium 6 fluorescent dye in INS-1 cells after 24 hour incubations performed under vehicle, IL-1β (5 ng/ml), NaB (0.5 mM), and IL-1β + NaB (5 ng/ml and 0.5 mM, respectively) treatment conditions. (**A)** The assay paradigm is illustrated. (**B)** Representative experimental raw fluorescence tracings. (**C)** Representative experimental baseline-normalized fluorescence tracings. (**D)** Quantification of baseline-normalized Ca^2+^ release from the ER. (**E)** Quantification of baseline-normalized SOCE. The values obtained in 8 independent experiments are represented by open circles, and the results are shown as the mean ± S.E.M; \*\*\**P* < .001, **** *P* < .0001.

### 3.4 Inhibition of HDAC improves cell viability

Impairments in SOCE have been linked with reduced ER Ca^2+^ storage and increased β cell death (6). To test whether HDAC inhibition with NaB improves β cell viability, INS-1 cells were exposed for 24 hours to IL-1β in the absence or presence of NaB, and cell viability was determined. IL-1β treatment significantly reduced cell viability; however, co-treatment with NaB significantly increased viability (Figure 4A). Consistent with this observation, NaB reduced the production of cleaved caspase-3 in IL-1β-treated INS-1 cells (Figure 4B-C).

**FIGURE 4.**
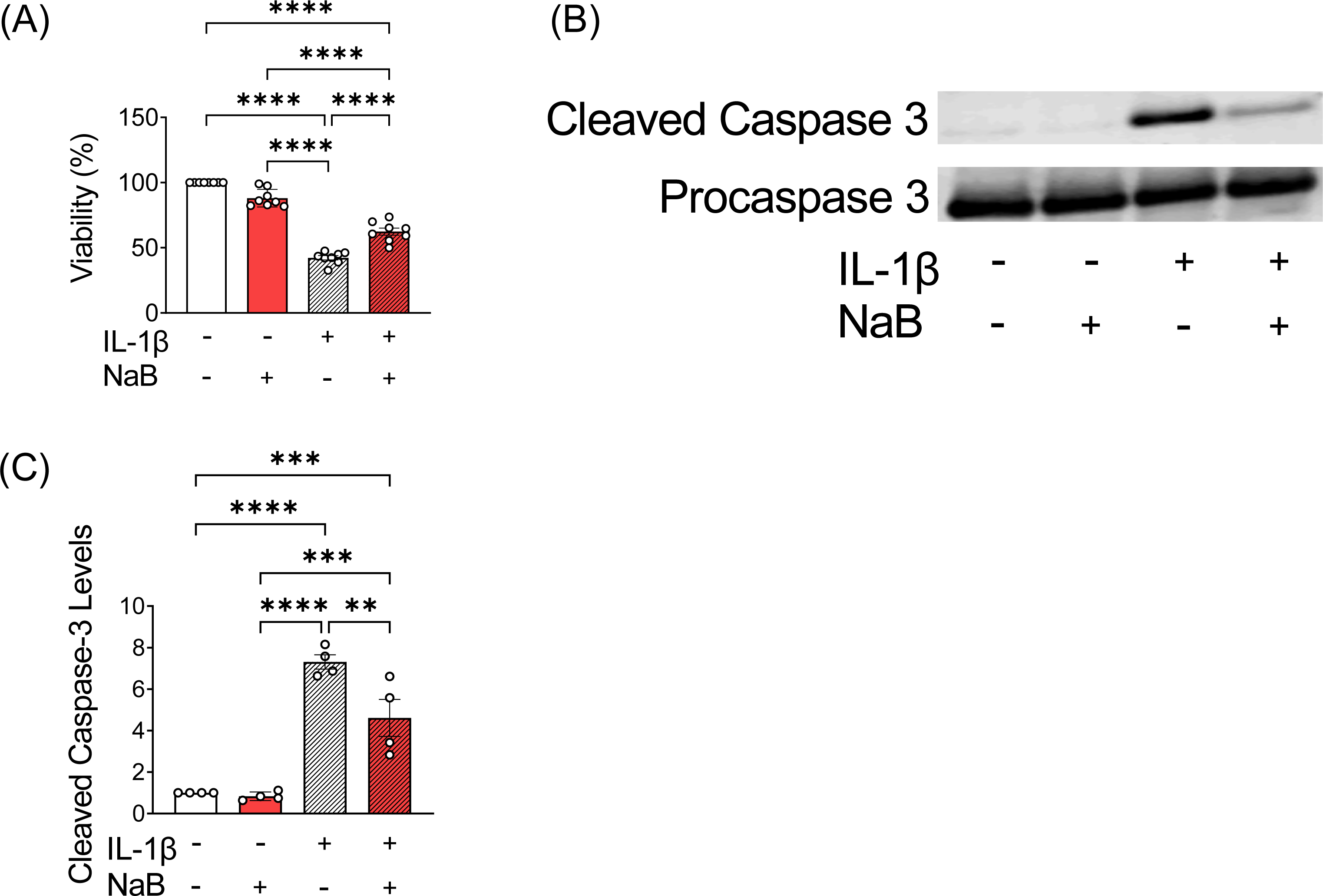
Inhibition of HDAC improves cell viability. (**A-C**) Cells were incubated in the absence or presence of IL-1β and NaB for 24 hours prior to assays of cell viability and caspase-3 cleavage. (**A)** CellTiter-Glo luminescent cell viability assay. n=8. (**B)** Immunoblot of cleaved caspase-3 and procaspase-3; n=4. (**C)** Quantification of relative signals within (**B**) for the ratio of cleaved caspase-3 normalized to procaspase-3. The results are shown as the mean ± S.E.M; \*\**P* < .01, \*\*\**P* < .001, \*\*\*\**P* < .0001.

### 3.5 NaB mediates rescued expression of SOCE-related genes

NaB exerts genomic effects through increasing histone acetylation and has been linked with increased acetylation in non-histone proteins, such as SP1 (24). Furthermore, NaB regulates metabolic and inflammatory genes and signaling pathways (25, 26). Temporal studies were performed to characterize the mechanisms underlying NaB-mediated improvements in SOCE response in INS-1 cells. Chronic (24 hours), but not acute (4 or 8 hours), NaB treatment improved SOCE in INS-1 cells exposed to IL-1β (Figure 5A). This finding raised the possibility that the beneficial effects of NaB involved changes in protein expression, which usually require time to manifest. Therefore, quantitative RT-PCR (qRT-PCR) was performed to measure the expression level of genes known to regulate SOCE. NaB treatment upregulated the expression of *Stim1*, *Stim2*, and *Orai1* mRNA in IL-1β-treated cells (Figure 5B). However, only the expression of *Stim1* was also reduced by IL-1β (Figure 5B), suggesting that NaB-mediated upregulation of *Stim1* expression could benefit SOCE and overall β cell function. The expression of *Orai2* was unchanged by the treatment conditions, while *Orai3* expression was increased in IL-1β-treated INS-1 cells and decreased upon co-treatment with IL-1β and NaB (Figure 5B). Similar to what we observed in INS-1 cells, cytokines reduced *Stim1* expression in mouse islets, while co-treatment with NaB rescued *Stim1* expression (Figure 5C). The expression of *Stim2*, *Orai1,* and *Orai3* did not vary among islet treatment groups, while *Orai2* expression levels were augmented by NaB treatment (Figure 5C).

**FIGURE 5.**
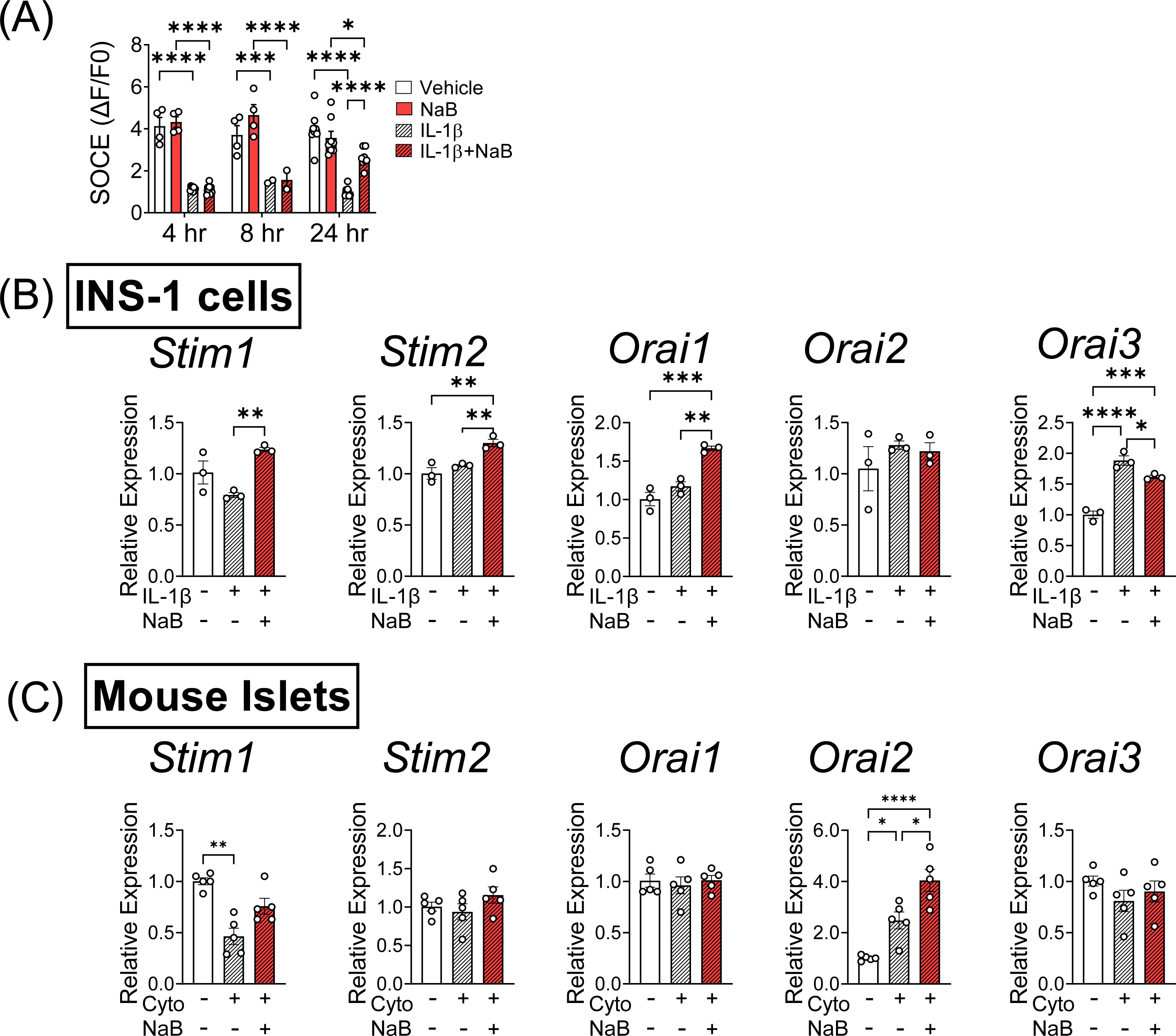
NaB treatment rescues the expression of SOCE-related genes under cytokine stress. (**A-B**) INS-1 cells were incubated in the absence or presence of IL-1β and NaB over a time course of 4-24 hours followed by Ca^2+^ imaging and analysis as described in Figure 3. Samples were also collected at 24 hours for quantitative RT-PCR analysis of gene expression levels. (**A)** Baseline-normalized quantitation of the SOCE response (ΔF/F0) from imaging tracings performed at 4, 8 and 24 hours of co-incubation. (**B)** Expression levels of SOCE-related genes (*Stim1*, *Stim2*, *Orai1*, *Orai2*, and *Orai3*). The expression level of β-actin served as an internal control for normalization of SOCE gene expression; n=3. (**C**) Mouse islets were incubated in the presence of cytokine mix and NaB for 24 hours and were subjected to quantitative RT-PCR analysis of genes related to SOCE, including *Stim1*, *Stim2*, *Orai1*, *Orai2*, and *Orai3*. The expression level of β-actin served as an internal control for normalization of SOCE gene expression and data are expressed relative to vehicle incubation; n=5. Cyto, cytokine. The results are shown as the mean ± S.E.M; \**P* < .05, \*\**P* < .01, \*\*\**P*<.001, \*\*\*\**P* < .0001.

### 3.6 NaB-mediated rescue of SOCE response is dependent upon the presence of STIM1

Consistent with the mRNA expression results reported above, increased STIM1 protein levels were detected in lysates prepared from INS-1 cells co-incubated with NaB and IL-1β (Figure 6A-B). Next, to determine if NaB improves SOCE under inflammatory conditions in a STIM1-dependent manner, we performed co-incubation studies in wildtype (WT) and STIM1 knockout (KO) INS-1 cells. Consistent with our hypothesis, NaB restored SOCE in IL-1β-treated WT INS-1 cells but failed to exert this effect in STIM1 KO cells (Figure 6C).

**FIGURE 6.**
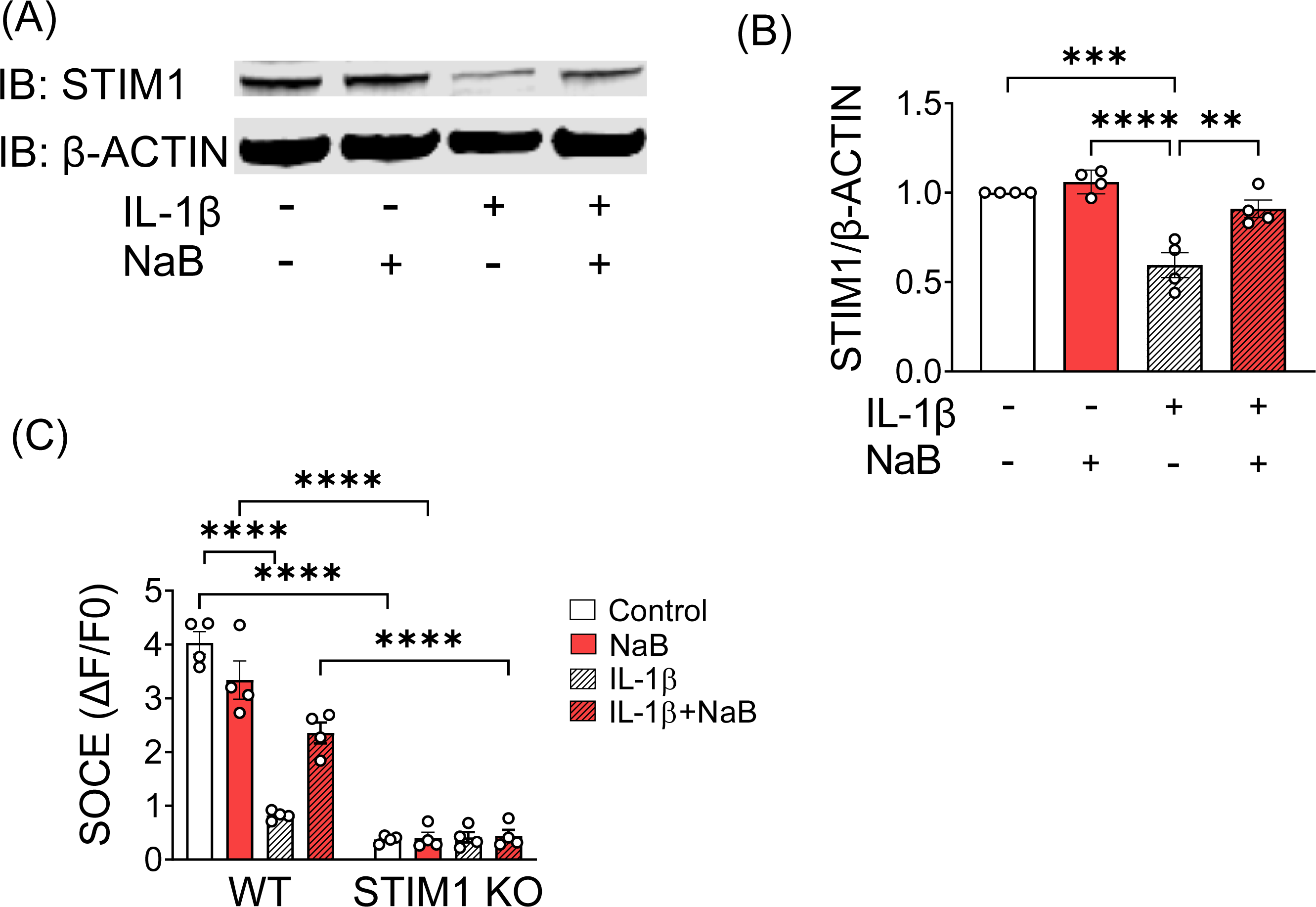
NaB-mediated rescue of SOCE response is dependent upon the presence of STIM1. (**A-B**) INS-1 cells were incubated in the absence or presence of IL-1β and NaB. Samples were collected at 24 hours for immunoblotting. (**A)** A representative immunoblot of STIM1 protein levels. The level of β-actin protein served as an internal control. (**B)** Quantitation of the ratio of STIM1 to β-actin protein levels for (**A**). Data are expressed relative to the vehicle group; n=4. (**C)** Wildtype (WT) and STIM1 KO INS-1 cells were incubated for 24 hours under the indicated conditions prior to imaging of SOCE. SOCE responses were calculated from experimental Ca^2+^ imaging tracings; n=4. The results are shown as the mean ± S.E.M; \*\**P* < .01, \*\*\**P* < .001, \*\*\*\**P* < .0001.

### 3.7 NaB rescues SOCE responses in cytokine-stressed β cells via the AKT-GSK-3 pathway

NaB has been shown to activate pancreatic AKT signaling in streptozotocin-treated rats, and AKT signaling regulates diverse cellular functions, including apoptosis, metabolism, and cell proliferation (27, 28). Therefore, we assayed changes in signaling that contributed to NaB-mediated effects on STIM1 and SOCE. Isolated mouse islets were treated with cytokines (IL-1β, IFN-γ, and TNF-α) in the presence or absence of NaB for 16 hours, and a phosphorylation array was used to screen for regulation of select proteins in the MAPK signaling pathway. The raw data of phosphorylated protein intensity are shown in Supplemental Figure 1, and the quantification is summarized in Figure 7. Exposure of islets to cytokines significantly decreased phosphorylation of GSK-3α, GSK-3β, and MKK6, while NaB co-treatment upregulated the phosphorylation levels of these proteins. Furthermore, the phosphorylation levels of AKT, ERK1/2, and CREB were reduced by cytokine treatment and increased by NaB co-treatment (Figure 7), suggesting the involvement of these phosphoproteins in NaB-mediated benefits.

**FIGURE 7.**
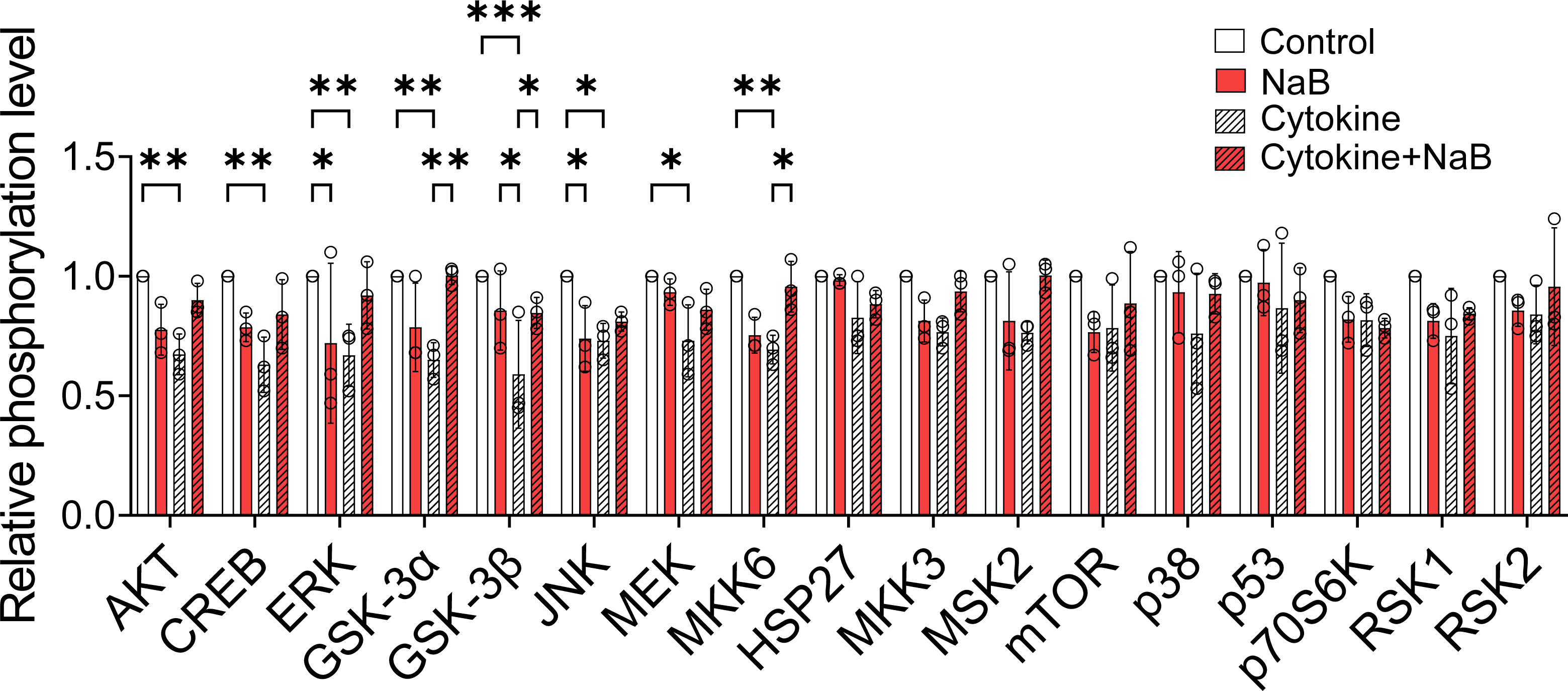
NaB rescues SOCE in cytokine-stressed β cells via the AKT-GSK-3 pathway. Signal transduction via the MAPK pathway was evaluated in WT C57BL/6J mouse islets after treatment with cytokine mix (5 ng/mL IL-1β, 100 ng/mL IFN-γ, and 10 ng/mL TNF-α) in the presence or absence of NaB for 16 hours. The islets were lysed and subjected to phosphorylation array procedures. The signals for every protein target were quantified and normalized to the positive controls in each membrane. For each protein target, the levels observed in the experimental group samples were expressed relative to the vehicle group levels. The values obtained in three independent experiments are represented by open circles, and the results are shown as the mean ± S.E.M; \**P* < .05, \*\**P* < .01, \*\*\**P* < .001.

## Discussion

The present study was designed to determine the effects and mechanisms of NaB, an HDI, in pancreatic β cells under proinflammatory stress. We found that NaB augments insulin secretion in human islets from donors with T2D. Furthermore, co-treatment of INS-1 cells or mouse islets with cytokines and NaB improves SOCE responses and cell viability, partially rescues insulin secretion, and improves Ca^2+^ oscillations. Mechanistic studies showed that NaB mediates its effects through the maintenance of *Stim1* expression and modulation of the AKT-GSK-3 signaling pathway. Importantly, STIM1 deletion in INS-1 cells abolishes NaB-mediated improvements in β cell function. Taken together, these results suggest a novel pathway through which NaB exerts protective metabolic effects in the diabetic β cell.

We and others have shown that loss of STIM1 in β cells impairs SOCE and results in a number of detrimental effects including reduced insulin secretion in response to glucose and fasiglifam, impaired glucose-stimulated Ca^2+^ oscillations, decreased ER Ca^2+^ levels, increased ER stress, reduced cell viability, and decreased β cell identity (6, 7, 14). Conversely, overexpression of STIM1 in human islets from donors with T2D improves insulin secretion, supporting the notion that STIM1 and SOCE play important roles in maintaining β cell health and function (6). However, the pathways leading to loss of STIM1 and reduced SOCE during diabetes have not been completely explored. Here, we show that the HDAC inhibitor, NaB, is able to improve β cell function under proinflammatory conditions in part by rescuing cytokine-mediated reductions in STIM1 expression and improving SOCE, Ca^2+^ oscillations, and insulin secretion.

NaB is a simple three carbon short-chain fatty acid. The primary dietary source of NaB is the host microbial fermentation of undigested dietary fiber, such as that found in legumes (29). The effects of NaB on cultured cells were initially recognized in studies characterizing proliferation, promotion of differentiation, and repression and induction of gene expression (30). Acetyl histone species accumulated in cultured mammalian cells when treated with NaB, and NaB was shown to suppress histone deacetylase activity *in vivo* and *in vitro* to promote histone acetylation (25, 31). The HDAC inhibition activity of NaB was confirmed in HEK293 cells, HMEC-1 cells (endothelial cells), L6 myotubes (skeletal muscle cells), and MAC-Ts cells (bovine mammary epithelial cells) (24). Histone hyperacetylation induced by NaB occurs through HDAC1, 2, and 3 and is estimated to affect the expression of a modest subset of ∼2% of mammalian genes (24). NaB changes gene expression through genetic mechanisms including regulation of gene chromatin accessibility, expression and/or modulation of signaling mechanisms that control transcription, or changes in levels of transcription factors and their complexes.

NaB has been linked with a number of anti-inflammatory effects in multiple cell types and disease states (25). NaB inhibited inflammation through suppression of NF-κB activation in patients with Crohn’s Disease (32), and is currently under clinical investigation for co-treatment along with probiotics for irritable bowel disease (IBD) (ClinicalTrials.gov, Identifier: NCT05013060) and pediatric obesity (33, 34), and as an add-on therapy to maintain remission in patients with ulcerative colitis (35). In endothelial cells, NaB reduced pro-inflammatory cytokine production in response to lipopolysaccharide (LPS) (36) and decreased activation of the <u>n</u>ucleotide-binding oligomerization domain-like receptors family <u>p</u>yrin domain containing <u>3</u> (NLRP3) inflammasome in adipocytes (25). Interestingly, NaB was also reported to trigger apoptosis in colon cancer cells and nasopharyngeal cancer cells in a SOCE-dependent manner (26). Of direct relevance to our study, NaB improves glycemia in streptozotocin and streptozotocin/high-fat diet treated mice (37). Mechanistically, these effects have been linked with reduced serum IL-1β levels, decreased NF-kB signaling in the total pancreas, decreased β cell apoptosis, and reduced islet ER stress (37).

These findings led us to explore whether NaB impacts pancreatic β cell health, function, and Ca^2+^ homeostasis. In the present study, NaB countered IL-1β-mediated impairment of INS-1 cell health and function in our measures of viability, active caspase, and secretory function. These results are consistent with recent findings that NaB improves β cell function and proliferation in mouse islets, INS-1E cells, and human EndoC-βH1 cells (38). Our discovery that NaB improves SOCE in INS-1 cells under proinflammatory stress is novel, and we extended our analysis to show that NaB mediates functional improvements in Ca^2+^ oscillation signals in response to glucose stimulation in mouse islets. In our studies, cytokine treatment of mouse islets abolished Ca^2+^ oscillations, while first and second phase oscillatory peaks partially returned with NaB co-incubation. Furthermore, we found that NaB improved GSIS from cadaveric human islets isolated from individuals with T2D.

In addition, using a phosphorylation array, we identified changes in AKT-GSK3 signaling in response to NaB treatment in β cells. These findings were notable since NaB was reported to enhance pancreatic AKT signaling in streptozotocin-treated rats (27), and AKT signaling regulates diverse cellular functions, including apoptosis, metabolism, and cell proliferation (39). Our findings extend these observations and implicate that changes in GSK-3 signaling contribute to NaB effects.

Overall, our data indicate that the HDAC inhibitor NaB promotes β cell health and function under proinflammatory conditions through STIM1-mediated control of SOCE and AKT-mediated inhibition of GSK-3. Further studies will determine whether NaB treatment could be a viable therapeutic strategy for the treatment of diabetes.

## Data Availability Statement

The data supporting the findings in this study are available in the Methods and/or Supplementary Material of this article. Further information and requests for resources and reagents should be directed to and will be fulfilled by the corresponding author. The study did not generate new unique reagents.

## Author Contributions

C.L. contributed to the conception and design of the study, data analysis and interpretation, collection and assembly of data, and manuscript writing. T.K. directed the study conception and design, contributed to acquisition, analysis, interpretation of data, and provided critical revision of the manuscript. F.S. contributed to the conception and design of the study and revision of the manuscript. S.A.W. contributed to the data collection and revision of the manuscript. P.S. contributed to data collection and revision of the manuscript. W.W. contributed to data collection and analysis. G.C. contributed data analysis. J.L. contributed to the conception of data analysis. M.S.R. contributed to data analysis. C.E.-M. directed funding acquisition, study conception and design, participated in the collection and assembly of data, contributed to data analysis, directed manuscript writing, and gave final approval of the manuscript. C.E.-M. is the guarantor of this work and, as such, has full access to all the data in the study and takes responsibility for the integrity of the data and the accuracy of the data analysis. All authors gave final approval of the manuscript.

## Nonstandard abbreviations

AKT: Ak strain transforming
Cyto: cytokine
Dz: diazoxide
ER: endoplasmic reticulum
ERK1/2: extracellular signal-regulated kinase 1/2
Fura-2-AM: Fura-2-acetoxymethyl ester
GSIS: glucose-stimulated insulin secretion
GSK-3: glycogen synthase kinase-3
HDAC: histone deacetylase
HDI: histone deacetylase inhibitor
IBD: irritable bowel disease
IL-1β: interleukin-1β
IFN-γ: interferon-γ
IP3: inositol triphosphate
KO: knockout
LPS: lipopolysaccharide
NaB: sodium butyrate
NLRP3: nucleotide-binding oligomerization domain-like receptors family pyrin domain containing 3
ROI: region of interest
RRID: Research Resource Identifiers
SAB: secretion assay buffer
SERCA: sarco-ER Ca^2+^ ATPase
SOCE: store-operated calcium entry
SSA: singular spectrum analysis
STIM1: stromal interaction molecule 1
STIM1 KO: STIM1 knockout cells
T1D: type 1 diabetes
T2D: type 2 diabetes
TG: thapsigargin
TNF-α: tumor necrosis factor-α
WT: wild type

## Acknowledgements

This work was supported by National Institute of Diabetes and Digestive and Kidney Diseases grants R01DK093954, R01DK127236, U01DK127786, R01DK127308, UC4DK104166 (to C.E.-M.), U.S. Department of Veterans Affairs Merit Award I01BX001733 (to C.E.-M.), and gifts from the Sigma Beta Sorority, the Ball Brothers Foundation, the George and Frances Ball Foundation (to C.E.-M.). W.W. was supported by the grant U24DK097771 from the NIDDK Information Network’s (dkNET) New Investigator Pilot Program in Bioinformatics. The authors acknowledge the support of the Islet and Physiology Core of the Indiana Diabetes Research Center (P30-DK-097512). The authors thank Dr. Emily Anderson-Baucum for her helpful advice and edits.

## Supplemental Figure Legends

**Supplemental Figure 1.**
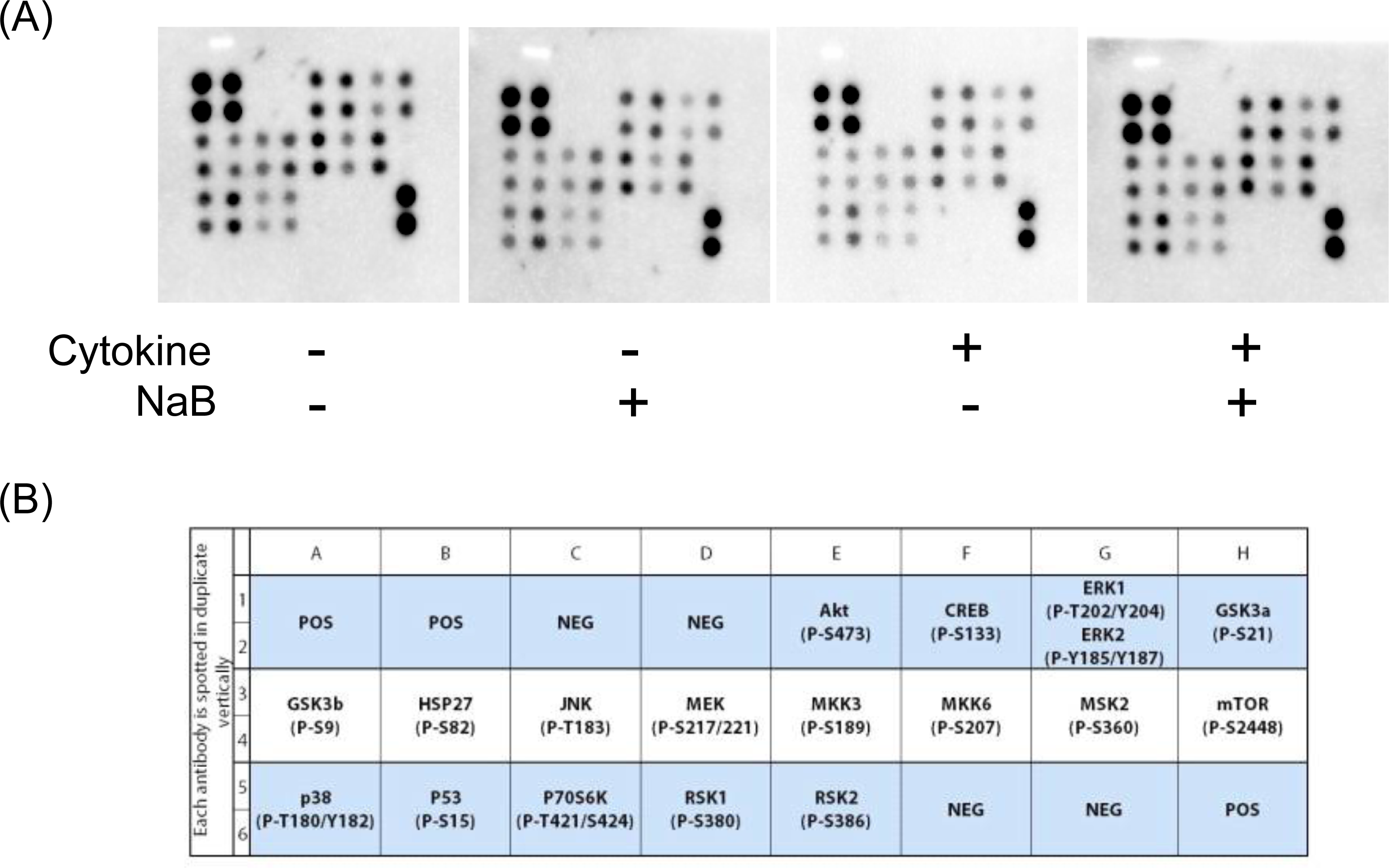
Representative membranes of phosphorylation array assay in mouse islets treated with cytokines in the presence or absence of NaB. Isolated mouse islets were treated with NaB and/or cytokines (IL-1β, IFN-γ, and TNF-α) for 16 hours. Cell lysates were incubated with membranes that were dotted with select phosphoprotein targets in the MAPK signaling pathway. (**A**) Membranes were imaged using chemiluminescence with ChemiDoc Imaging System. (**B**) Table of phosphoprotein targets and their location on the membrane.

